# The emergence of multiple retinal cell types through efficient coding of natural movies

**DOI:** 10.1101/458737

**Authors:** Samuel A. Ocko, Jack Lindsey, Surya Ganguli, Stephane Deny

**Affiliations:** Department of Applied Physics, Stanford and Google Brain, Mountain View, CA

## Abstract

One of the most striking aspects of early visual processing in the retina is the immediate parcellation of visual information into multiple parallel pathways, formed by different retinal ganglion cell types each tiling the entire visual field. Existing theories of efficient coding have been unable to account for the functional advantages of such cell-type diversity in encoding natural scenes. Here we go beyond previous theories to analyze how a simple linear retinal encoding model with different convolutional cell types efficiently encodes naturalistic spatiotemporal movies given a fixed firing rate budget. We find that optimizing the receptive fields and cell densities of two cell types makes them match the properties of the two main cell types in the primate retina, midget and parasol cells, in terms of spatial and temporal sensitivity, cell spacing, and their relative ratio. Moreover, our theory gives a precise account of how the ratio of midget to parasol cells decreases with retinal eccentricity. Also, we train a nonlinear encoding model with a rectifying nonlinearity to efficiently encode naturalistic movies, and again find emergent receptive fields resembling those of midget and parasol cells that are now further subdivided into ON and OFF types. Thus our work provides a theoretical justification, based on the efficient coding of natural movies, for the existence of the four most dominant cell types in the primate retina that together comprise 70% of all ganglion cells.

## 1 Introduction

The time honored principle that the visual system evolved to efficiently encode the structure of our visual world opens up the tantalizing possibility that we can predict, *ab initio*, the functional organization of visual circuitry simply in terms of the statistical structure of natural scenes. Indeed, efficient coding theory has achieved several successes in the retina by simply considering coding of static spatial scenes [1, 2, 3] or mostly temporal sequences [4]. However, such theories have not yet accounted for one of the most salient aspects of retinal computation, namely the existence of a diversity of retinal ganglion cell types, each forming a mosaic that uniformly tiles the visual field [5].

A few theoretical studies have suggested reasons for different cell types. One suggestion is the feature detector hypothesis [6, 7, 8], or the need to detect highly specialized, behaviorally relevant environmental cues. However, many cell types respond broadly to general classes of stimuli whose *direct* behavioral relevance remains unclear [9, 10, 11]. Another line of argument involves metabolic efficiency. In particular the division of ganglion cells into rectifying ON and OFF populations is more metabolically efficient than linear encoding with a single population [12], and the asymmetry between ON and OFF cells can be related to the asymmetric distribution of light intensity in natural spatial scenes [13]. Another efficient coding argument explains why two populations with similar receptive fields (RFs) have different activation thresholds in the salamander retina [14].

Here we go beyond previous efficient coding theories of the retina by optimizing convolutional retinal models with multiple cell types of differing spatial densities to efficiently encode the *spatiotemporal* structure of natural *movies*, rather than simply the spatial structure in natural scenes. Indeed, psychophysical studies of human sensitivity [15, 16] suggest our visual system is optimized to process the spatiotemporal information content of natural movies. Our theory enables us to account for several detailed aspects of retinal function. In particular, the primate retina is dominated by four types of ganglion cells, ON midget and parasol cells and their OFF counterparts. Together, these types constitute 68% of all ganglion cells [17], and more than 95% in the central retina [18]. Midget cells are characterized by (1) a high density of cells (52% of the whole population), (2) a small spatial RF, (3) slow temporal filtering, and (4) low sensitivity – as measured by the slope of their contrast-response function. In contrast, parasol cells are characterized by (1) a low density of cells (16% of the whole population), (2) a large RF, (3) fast temporal filtering and (4) high sensitivity [19, 20, 21, 22]. Moreover, the density ratio of midget to parasol cells systematically decreases across retinal eccentricity from the fovea to the periphery [23].

Remarkably, our theory reveals how all these detailed retinal properties arise as a natural consequence of the statistical structure of natural movies and realistic energy constraints. In particular, our theory simultaneously accounts for: (1) why it is beneficial to have these multiple cell types in the first place, (2) why the four properties of cell density, spatial RF-size, temporal filtering speed, and contrast sensitivity co-vary the way they do across midget and parasol types, and (3) quantitatively explains the variation in midget to parasol density ratios over retinal eccentricities. Moreover, our theory, combined with simulations of efficient nonlinear encoding models, also accounts for the existence of both ON and OFF midget and parasol cells. Thus simply by extending efficient coding theory to multiple cell types and natural movies with a realistic energy constraint, we account for cell-type diversity that captures 70% of all ganglion cells.

## 2 A theoretical framework for optimal retinal function

### Retinal model

We define a ganglion cell type as a convolutional array of neurons sampling linearly from a regularly spaced array of N_p_ photoreceptors, indexed by *i* = 0,…, N_p_ – 1 (Fig. 1A). We model photoreceptors as linearly encoding local image contrast. We also assume the N_*C*_ ganglion cells of cell type *C*, indexed by *j* = 0,…, N_*C*_ – 1, each have a common spatiotemporal RF, defined as F_*C*_(*i*, Δ*t*), whose center is shifted to a position *j* · s_*C*_ on the photoreceptor array. Here *s_C_* is the *convolutional stride* of type *C*, which is an integer denoting the number of photoreceptors separating adjacent RF centers of ganglion cells of type *C*, so that N_p_ = *s_C_*N_*C*_. This yields a retinal model

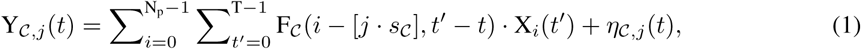

where X_*i*_(*t*′) is activation of photoreceptor *i* and time *t*′, *η_c,j_*(*t*) is additive noise that is white across both space (*C* and *j*) and time (*t*) with constant variance 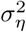 [24], and Y*_Cj_*(*t*) is the firing rate of ganglion cell *j* of type *C*. We work in one spatial dimension (generalizing to two is straightforward). This linear model will provide conceptual insight into spatiotemporal RFs of different cell types through exact mathematical analysis. However, in Sec. 5 we also consider a nonlinear version of the model to account for rectifying properties of different cell types.

**Figure 1:**
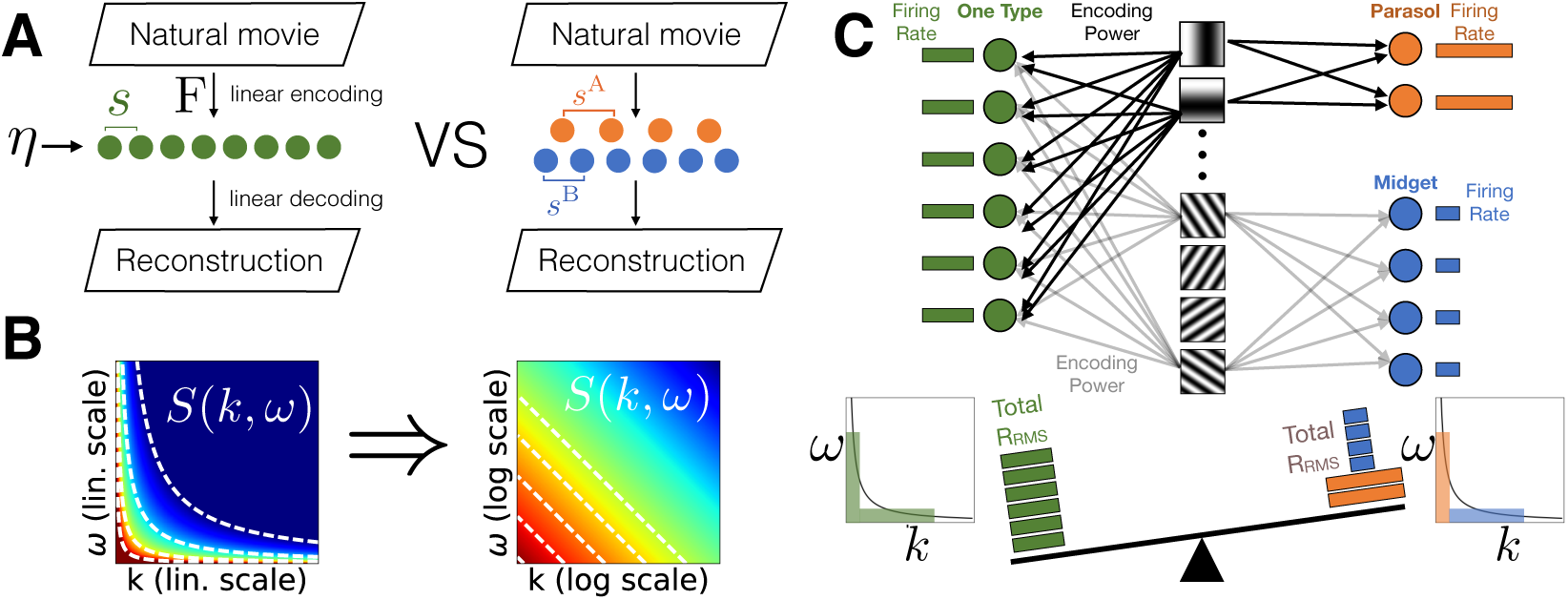
An efficient coding model for multiple convolutional cell types. A) Natural movies are encoded in a set of ganglion cells (circles) through linear filters corrupted by noise (Eq. 1). Left: one convolutional cell type with stride s (green). Right: two different cell types (blue and orange) with different strides *s*^𝔸^ and *s*^𝔹^. B) The Fourier power spectrum S(*k*, *ω*) of natural movies decays as a separable power law in both spatial (*k*) and temporal (*ω*) frequency (see Eq. 2). Dashed lines are iso-contours of constant power in linear (left) and logarithmic (right) axes with power varying from high (red) to low (blue). Note that, aside from the origin, the most powerful Fourier modes are contained in two distinct regions: (1) low *k*, high *ω*, and (2) high *k*, low *ω*. C) We will show that two convolutional cell types (orange, blue) encode visual information more efficiently than one type (green) by specializing their filters to cover these two regions. The orange, parasol-like cell type specializes to region (1) using a small number of cells at large stride with large spatial RFs (low *k*), and fast temporal filters (high *ω*) that fire sensitively at high rates. The blue, midget-like cell type specializes to region (2) using a large number of cells at small stride with small spatial RFs (high *k*) and slow temporal filters (low *ω*) using low firing rates. Together, these two specialized cell types can encode photoreceptor input patterns with the same fidelity as a single, undifferentiated cell type (green), with less total firing rate.

### Natural movie statistics

We approximate natural movies by their second order statistics (Fig. 1B), assuming Gaussianity. Natural movies are statistically translation invariant, and so their second order statistics are described by their Fourier power spectrum, which follows an approximately space-time separable power law as function of spatial (*k*) and temporal (*ω*) frequency [25, 26, 27]:

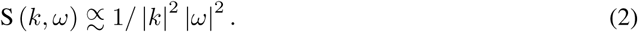

### Optimization Framework

We assume the objective of the retina is to faithfully encode natural movies while minimizing overall ganglion cell firing rate. We quantify encoding fidelity by the amount of input variance *V* explained by the optimal minimum mean squared error (MMSE) reconstruction of photoreceptor activity patterns from ganglion cell outputs by a linear decoder. Moreover, we assume an overall penalty on output firing rate that is proportional to a power *p* of the rate. This yields an objective function to be maximized over the set of convolutional filters {F_*C*_ }:

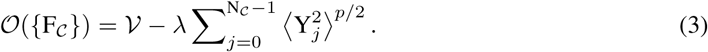

Here λ is a parameter that trades off between the competing desiderata of maximizing encoding fidelity versus minimizing firing rate. Our final results will focus on the choice *p* =1, motivated by the linear relationship between metabolic cost and firing rate [28, 29]. This choice is similar to an 𝓵_1_ penalty used in [30, 31, 2]. However we also consider more general *p*, including *p* = 2, both to connect to prior work on efficient coding [1,3] and as a building block for solving the *p* =1 case.

### Outline

In Sec. 3, we prove mathematically that multiple cell types enable a more efficient code (i.e. same or better encoding fidelity with lower firing rates) than a single cell type, as long as *p* < 2. The fundamental idea is that different cell types allow higher efficiency by *specializing* to different regions of the power spectrum of natural movies (Fig. 1C). In Sec. 4, we find the optimal cell types for natural movies, demonstrating that the best two-type strategy substantially out-performs the best one-type strategy. We then compare these optimal types to midget and parasol cells in the primate retina and find striking agreement between optimal and biological cell types. In Sec. 5, we extend our theory to non-linear ganglion cells and account for both ON and OFF midget and parasol cells.

## 3 Mathematical proof of the benefit of multiple cell types

Here we derive mathematically how multiple specialized cell types can confer an efficient coding advantage compared to a single cell type (Fig. 1A,C). In Sec. 3.1, we start with the simple case of a single cell type with stride 1 yielding an equal number of ganglion cells and photoreceptors, encoding static images [1, 32]. We then extend this framework to varying strides (Sec. 3.2), encoding natural movies (Sec. 3.3), and multiple cell types, proving that they can confer an advantage (Sec. 3.4). We will solve for the optimal RFs and strides of each cell type in Sec. 4.

### 3.1 Encoding N_p_ photoreceptors with N_C_ = N_p_ ganglion cells of a single type

As a warmup, for purely spatial scenes, we first consider optimizing the single cell-type retinal filter F_C_ (*i*, Δ*t*) in Eq. 1 under the objective function in Eq. 3, in the simple case of stride *s_C_* = 1 so that N_C_ = N_p_. Since we only have one cell type, we drop the cell-type index *C* in the following. The case of N_C_ = N_p_ simplifies because we can ignore aliasing [33], which we address in the next section. Thus we can show (App. A) that each spatial Fourier mode 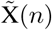 of photoreceptor patterns maps in one-to-one fashion onto a single spatial Fourier mode 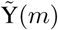 of ganglion cell patterns (Fig. 2A1):

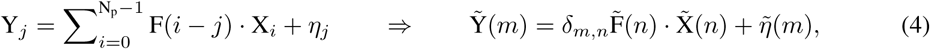

where *δ* is the Kronecker delta function, *n* ∊ {–N_p_/2 + 1,… 0,1,… N_p_/2} indexes photoreceptor Fourier modes, *m* ∊ {—N_C_/2 + 1,… 0,1,… N_C_/2} indexes ganglion cell Fourier modes, and 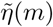 is the spatial Fourier transform of the noise (which also has variance 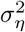). 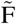 is the Fourier transform of F across photoreceptors, rescaled by 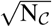. Each mode number *n* (*m*) corresponds to a photoreceptor (ganglion cell) spatial frequency k_*n*_ ≡ 2*πn*/N_p_ (*p_m_* ≡ 2*πm*/N_C_). The power S(*n*) in photoreceptor mode *n* is simply proportional to the power S (*k*, *ω*) in natural movies (Eq. 2) evaluated at spatial frequency *k* = *k*_|*n*|_. Finally, because image statistics are translation invariant, the objective (Eq. 3) can be written (App. A) in terms of independent photoreceptor spatial modes (here *p* = 2):

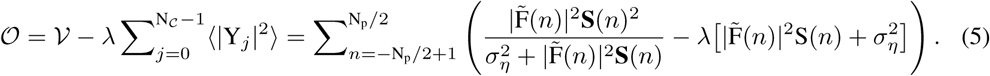

Thus 𝓞 can be maximized independently for each filter mode *n*, yielding the optimal filter (App. A):

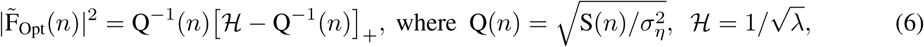

where Q(*n*) is a measure of the *quality* of photoreceptor Fourier mode, or input channel *n*.

**Figure 2:**
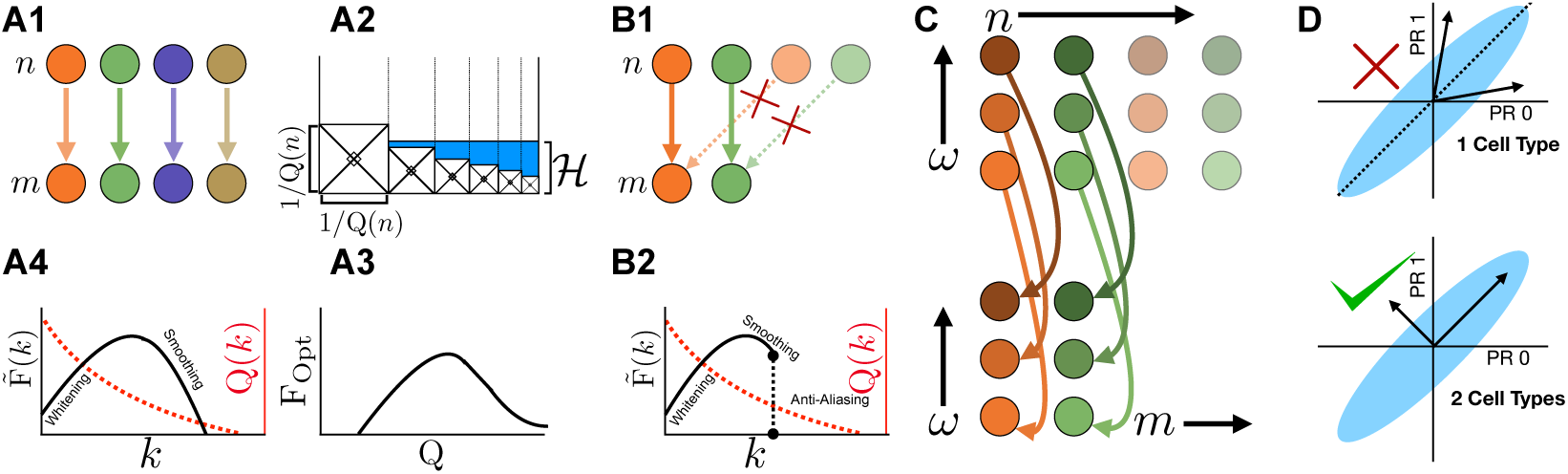
A1) In the convolutional framework of [1], every photoreceptor spatial frequency (upper dots) maps to a corresponding ganglion cell spatial frequency (lower dots). A2) The optimal filter strength assigned to each mode *n* can be viewed as the volume of water assigned to a corresponding beaker whose base height and width are inversely related to channel quality Q(*n*), and the water height 𝓗 is inversely related to the firing rate penalty parameter λ. A3) The optimal filter strength is a non-monotonic function of channel quality. A4) This also leads to non-monotonic optimal filter strength as a function of spatial frequency. B1) With fewer ganglion cells than photoreceptors, the optimal filter will sample *only* from the lowest photoreceptor spatial frequencies that map in one-to- one fashion to the lowest ganglion cell spatial frequencies, and will ignore the higher photoreceptor frequencies that alias to the same ganglion cell frequencies (App. B). B2) Thus the optimal filter achieves a similar solution as in A4, with an additional upper bound on frequency to avoid aliasing. C) In spacetime, the optimal filter maps spatiotemporal photoreceptor frequencies (upper dots) to ganglion cell frequencies (lower dots) in a one-to-one fashion, ignoring higher photoreceptor spatial frequencies to avoid spatial aliasing. Different spatial (temporal) frequencies are indicated by shade (color). D) (See Sec. 3.4) The blue ellipsoid illustrates a correlated stimulus covariance across two photoreceptors (PR). Top: the two arrows denote two filters 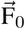 and 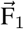 of two ganglion cells of a single cell type, related by a convolutional translation, modulo N_p_ = 2 and therefore reflection-symmetric about the diagonal (Sec. 3.4). Bottom: the two arrows denote two rotated filters 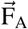 and 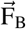 that each specialize to a different eigenbasis vector of the stimulus covariance, thereby differentiating into two cell types, enabling a more efficient neural code with the same fidelity but lower firing rate cost.

This solution has an appealing water-filling [34] interpretation (Fig. 2A2) in which each channel *n* of quality Q(*n*) corresponds to a beaker with base height and width both equal to Q(*n*)^−1^. These beakers are filled with water up to height 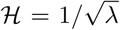, and the power 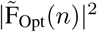 assigned to filter mode *n* is simply the *volume* of water in beaker *n*. Thus extremely low quality channels with beaker base Q(*n*)^−1^ greater than the water height 𝓗 are not used. Similarly, high quality channels with a low base are not assigned much filter strength because they are also narrow. Thus the optimal solution assigns filter strength as a non-monotonic function of channel quality (Fig. 2A3), favoring channels of intermediate quality, eschewing extremely low quality channels that do not contribute much to encoding fidelity, while attenuating channels that are already high-quality whose amplification would yield a cost in firing rate that outweighs the improved coding fidelity. As the penalty λ in firing rate is reduced, the water height 𝓗 increases, and more lower quality channels are used by the optimal filter.

Because the power spectrum of natural movies decays with spatial frequency [25], higher (lower) quality channels correspond to lower (higher) spatial frequencies. Thus the non-monotonic optimal filter strength as a function of channel quality (Fig. 2A3) leads to two qualitative effects (Fig. 2A4): (1) the attenuation of very low frequency high quality channels relative to intermediate frequency channels (spatial whitening) driven primarily by the need to lower firing rate, and (2) the eschewing of very high frequency low quality channels (spatial smoothing), which do not contribute strongly to encoding fidelity.

### 3.2 Encoding N_p_ photoreceptors with N_C_ < N_p_ ganglion cells of a single type

In the case of strides greater than 1, with fewer ganglion cells than photoreceptors, more than one spatial Fourier mode of photoreceptor activity can map to the same spatial Fourier mode of ganglion cell activity, a phenomenon known as *aliasing.* Indeed, not only does photoreceptor mode index *m* map to ganglion cell mode index *m*, as in Sec 3.2, but so does every other photoreceptor mode *n* separated from *m* by an integer multiple of N_C_ (Fig. 2 B, App. B), yielding the map

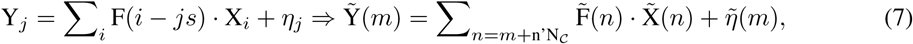

where n’ ranges over integers such that *n* = *m* + n’N_C_ enumerates all photoreceptor frequencies *n* within the bounds —N_p_/2 < n < N_p_/2 which alias to the same ganglion cell frequency *m*. Despite the many-to-one map from photoreceptor to ganglion cell Fourier modes, one can still optimize Eq. 3 independently over different filter modes akin to Eq. 5 through the following argument. The firing of each ganglion cell frequency *m* comes from the set of photoreceptor frequencies which alias to it, i.e. *n* = *m* + n’N_C_ (Fig. 2 B1). First we show that it is optimal for a single ganglion cell mode *n* to draw

*only* from the input eigen-mode with largest power (App. B.1). Now given the lowest photoreceptor spatial frequencies have the highest power, the optimal convolutional filter should sample only from the lowest photoreceptor frequencies, and not from the higher aliasing frequencies. Thus the optimal filter has 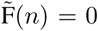 for all *n* with |*n*| > Nc/2. Therefore, the optimal filter in frequency space is simply a scaled, truncated version of Eq. 6 (Fig. 2 B1, B2):

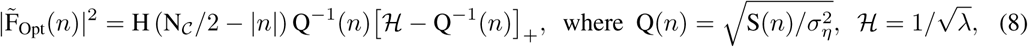

where H is the Heaviside function.

Note the optimal upper frequency cutoff to avoid aliasing naturally yields tiling, in which the spatial RF width becomes proportional to the stride [33]. To see this, consider a cell type with a stride in physical space of length s. Spatial frequencies higher than *O*(1/*s*) will lead to aliasing, yielding a frequency cut-off *k_s_* α 1/s. Further assume, for simplicity, that the water-filling solution fills all spatial frequency modes up to *k_s_* with the same amplitude, and chooses the same phase. This yields a box Fourier spectrum whose inverse spatial RF is a *sinc* function whose first zero crossing occurs at spatial scale 1/*k_s_* α s. Thus the RF width is proportional to stride, and cells with high (low) frequency cut-offs have small (large) RF widths and strides.

### 3.3 Generalizing the framework to spatiotemporal movies for a single cell type

With the addition of time, we can Fourier transform Eq. 1 in both space and time, yielding

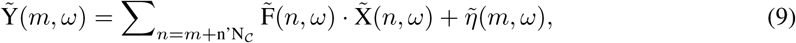

where X*_i_*(*t*′) is the activity of photoreceptor *i* at time *t*′, 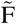 is the Fourier transform of F rescaled by 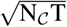 and 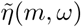 is the Fourier transform of the noise. Note that with fewer ganglion cells than photoreceptors, there will be a many-to-one map from photoreceptor spatial frequency to each ganglion cell spatial frequency as in Eq. 7, but a one-to-one map from photoreceptor to ganglion cell temporal frequencies (Fig. 2C, App. C). As in Sec. 3.2, the optimal filter will map the lowest photoreceptor spatial frequencies one-to-one to the lowest ganglion cell frequencies, while ignoring higher photoreceptor spatial frequencies to avoid aliasing. Moreover, within this aliasing constraint, spatiotemporal photoreceptor frequencies map one-to-one to ganglion cell frequencies, yielding an optimal solution given by Eq. 8 with channel quality depending on spacetime power S(k_*n*_, *w*).

### 3.4 Interplay of firing rate penalty and the benefit of multiple cell types

To build intuition for when and why multiple cell types can enable more efficient neural codes, we consider the simplest possible scenario of N_p_ = 2 photoreceptors and two convolution ganglion cells of a single type. The stimulus statistics and ganglion cell filters are given by (see also Fig. 2D):

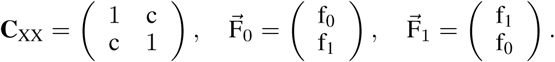

Note the two filters are equal up to a translation (modulo N_p_) and therefore obey the convolutional constraint. Let’s call D the optimal decoder. Then the reconstruction 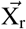 of the input is:

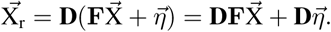

Here **F** is a 2 by 2 filter matrix whose rows are given by the two ganglion cell filters. Now the decoding performance is unaffected by an orthogonal rotation of the rows of **F**. Indeed, when **F** → **RF**, we can transform **D** → **DR**^−1^ yielding the reconstruction

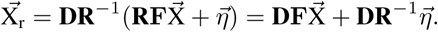

Because **R** is a rotation (i.e. **R**^-1^ = **R**^T^) and 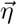 is isotropic Gaussian white noise, the statistics of 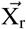 conditioned on 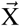, and thus the explained variance *V*, is unchanged by the rotation. More formally, the explained variance can be computed to be 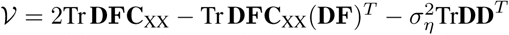 [35], and is independent of the transformation effected by **R**. This yields an entire manifold of optimal filter matrices **F** with the *same* explained variance *V*.

Now consider a *particular* choice of rotation **R** that rotates the two convolutional filters into the eigenbasis of **C**_XX_:

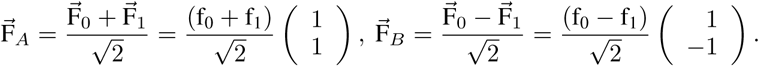

The rotated filters 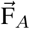 and 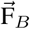 are no longer related by any translation. Thus the convolutional constraint is relaxed and they are analogous to two different cell types. We now compare the signal component of the total firing rate cost for the single cell-type convolutional filters, given by:

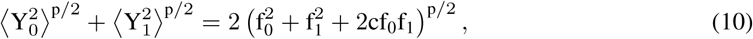

with the rotated, specialized, two-cell type filters, given by

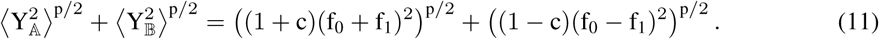

As long as c ≠ 0 and p < 2, the rotated (Eq. 11) two-type solution uses a lower firing rate budget than the one-type solution (Eq. 10). We generalize this proof in App. D to arbitrary numbers of cells, convolutional types and natural movie statistics. Thus intriguingly we find a sharp transition in the exponent *p* relating firing rate to cost, with multiple cell types favored if and only if *p* < 2.

Some prior work on efficient coding [1,3] employed an *𝓵*_2_ penalty on firing rate (i.e. *p* = 2), while others [30, 31, 2] have employed an *𝓵*_1_ penalty (i.e. *p* = 1 in our Gaussian scenario). We note that energetic considerations suggest that metabolic cost is linearly related to firing rate [28, 29] (i.e. *p* = 1). Prior knowledge that multiple retinal cell types do indeed exist, in addition to these energetic considerations, lead us to consider *p* = 1 in the following, corresponding to a penalty on the root-mean-squared (RMS) firing rates, summed over all cells.

## 4 Comparison of theoretically derived cell types to primate retinal cell types

Here we optimize the efficient coding objective in Eq. 3, with a metabolically motivated firing rate penalty corresponding to *p* = 1, using the procedure described in App. E. In particular, for the same total RMS firing rate budget (obtained by increasing λ in Eq. 3 until we first match this budget) we found both the optimal single (Fig. 3A) and two (Fig. 3B) cell type solutions. For the two-cell type solutions, we additionally scanned the two strides of each cell type. Given that for each cell type *C*, the number of cells N_C_ and stride *s_C_* are related by *s_C_*N_C_ = N_p_, varying the two strides is equivalent to varying the fractions of each cell type. All such two-cell type solutions had the same RMS rate, but varying encoding fidelity (the variance explained term *V* in Eq. 3). Thus for each firing rate budget, we find an optimal cell-type ratio with highest encoding fidelity. In Fig. 3AB we employed a budget that yielded an optimal cell-type ratio matching that of midget to parasol cells in the primate retina [18]. However, the general structure of the resultant RF power spectra in Fig. 3AB is robust to the choice of total RMS firing rate budget.

**Figure 3:**
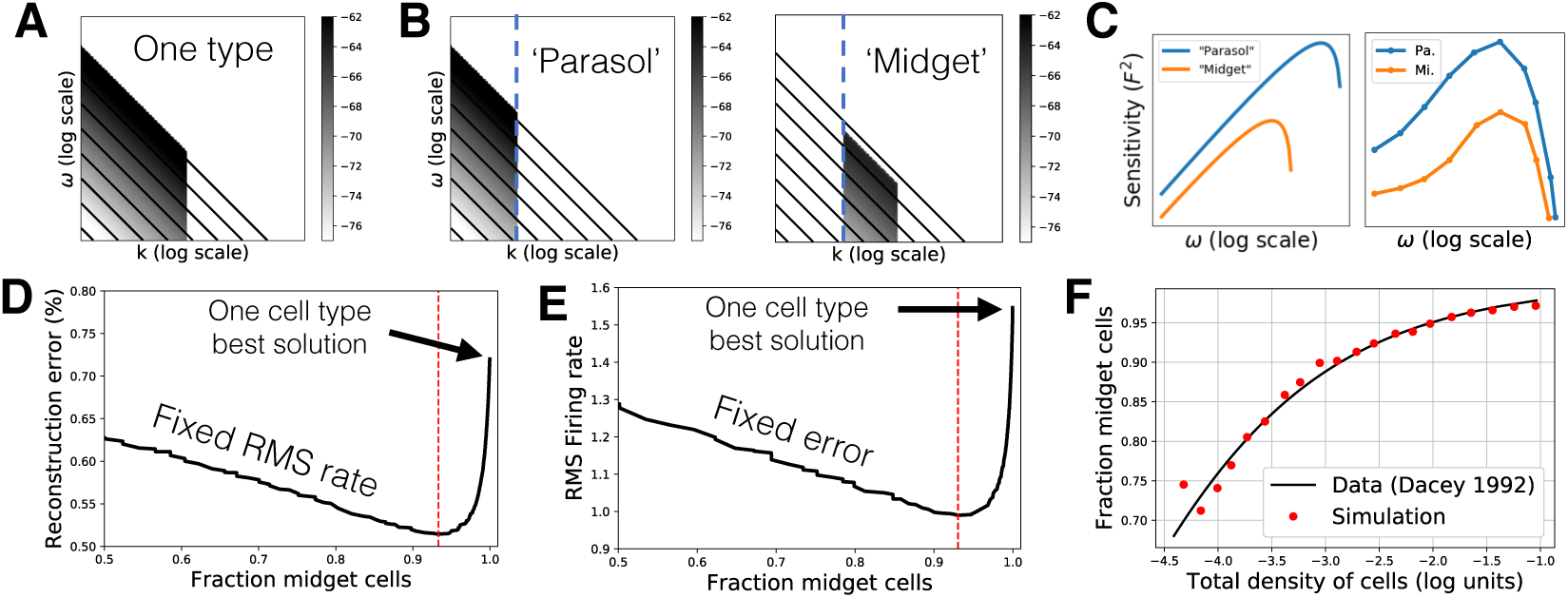
Optimal cell types match properties of midget and parasol cells. A) Optimal RF power spectra for a single cell type with darker shades denoting higher filter strength. Black lines are iso-contours of the power spectrum of natural movies. B) Optimal RF power spectra for two cell types with same conventions as A. C) Left: RF power spectrum of the two cell types along the frequency axis at fixed spatial frequency (the blue dashed lines in B). Right: Measured sensitivity or contrast gain as a function of temporal frequency of real midget and parasol cells (reprinted from [36]). D) Reconstruction error as a function of the fraction of midget cells for the RMS firing rate budget at which the optimal fraction is 93% (red line), consistent with the fraction found at a certain retinal eccentricity [18]. E) RMS firing rate budget as a function of the fraction of midget cells, for a fixed reconstruction error. Note the optimal one type solution (last point on the right) requires a 50% higher firing rate than the optimal two-type solution. F) Fraction of midget cells as a function of the total density of cells. Red points: optimal fractions predicted by theory (only one parameter was fitted to the data, see text). Line: fit to the fraction of midget across the human retina estimated from [23].

Remarkably, this general structure of the theoretically derived two-cell-type solution matches many properties of biologically observed primate retinal cell types. The first type corresponds to parasol cells, covering low spatial frequencies (implying large spatial RFs with large stride and low number density) and high temporal frequencies (implying fast temporal filtering). The second type corresponds to midget cells, covering a large number of high spatial frequencies (implying small spatial RFs with small stride and high number density) and low temporal frequencies (implying slow temporal filtering). Moreover, a single slice of the RF power spectrum along temporal frequency at a fixed spatial frequency (Fig. 3C, left) reveals that our theoretically derived “parasol” cell type has higher sensitivity (i.e. filter strength, or gain between input-contrast and output response) compared to the “midget” cell type, consistent with observations from the primate retina (Fig. 3C, right [36]). Thus as promised in Fig. 1C, the striking covariation of the four distinct features (cell density, spatial RF-size, temporal filtering speed, and contrast sensitivity) across the two dominant primate retinal cell types, arises as a simple emergent property of the two tailed structure of the natural movie power spectrum. By specializing to these two tails, the two-cell type solution in Fig. 3B can achieve *higher* encoding fidelity at the *same* RMS firing rate budget compared to the single-cell type solution in Fig. 3A. Indeed for the common firing rate budget chosen in Fig. 3AB, the optimal two-cell type solution achieves a 34% reduction in reconstruction error compared to the single type solution (Fig. 3D). Conversely, at a fixed reconstruction error (of 0.5%), two cell types are 33% more efficient than one in terms of total RMS firing rate (Fig. 3E). More generally, across for any non-zero firing rate budget the two-type solution achieves higher encoding fidelity, and for any desired encoding fidelity, the two-type solution requires lower firing rates.

The fixed budget shown in Fig. 3D and the fixed reconstruction error shown in Fig. 3E, were chosen such that optimal fractions of midget and parasol cells were 93% and 7%, respectively, consistent with those found at certain eccentricities of the primate retina [23]. However, in our model the optimal fractions change as the firing rate budget is increased (or equivalently, as the reconstruction error is decreased). The total density of cells in the optimal solution computed by the model also varies with the firing rate budget. Thus, our model makes a specific numerical prediction relating total cell density to the ratio of midget to parasol cells. The total density of cells varies across eccentricity by 3 orders of magnitude in the primate retina. In Fig. 3F, we plot the predicted evolution of the percentage of midget cells with cell density and compare it to the evolution of this percentage estimated from biological data [23] (see App. F for estimation method). Our model involves only one adjustable parameter to account for our arbitrary choice of units of cell density. Remarkably, we find an excellent match between theory and experiment in Fig. 3F, providing further evidence that the principle of efficient encoding of natural movies under a limited firing rate budget may be driving the functional organization of the primate retina.

## 5 A neural network simulation for linear-non-linear neurons

While the linear theory accounts for several properties of midget and parasol cells, it suffers from two main deficiencies. First, like previous efficient coding theories [1, 32], it only predicts the power of RF Fourier spectra, leaving the phase, and therefore the full spacetime RF unspecified. Second, it cannot account for rectifying nonlinearities, leading to the partition of ganglion cells into ON and OFF types. Here we remedy these deficiencies through neural network simulations, in which we nonlinearly autoencode natural movies with *two* spatial dimensions and one temporal dimension using three-dimensional convolutional neurons (full simulation details are given in App. G).

The main simulation ingredients include: (1) enforcing nonnegativity of neural firing rates through a ReLU nonlinearity in the ganglion cell encoding layer, (2) an 𝓵_2_ penalty on total weight magnitude, corresponding to a cost for synaptic connections [37], (3) encouraging decoding input stimuli with a short but non-zero temporal lag [38, 39], (4) implementing a firing rate budget with an 𝓵_1_ penalty on total firing rate. We assume four cell types, and optimize the number of cells allotted to each type. To match the fact that our image contrast distribution is zero mean Gaussian, and therefore symmetric about the origin, we pair the types into two pairs of equal-size populations and keep the number of cells the same across each pair during this optimization, expecting that ON and OFF homologous types will emerge. It would be interesting to explore skewed image statistics and test whether these would yield ON-OFF asymmetries, as are found in biological retinas [13]. We confirmed that our equal pairing of types is a locally optimal cell type allocation (for our symmetric image statistics) by performing a stability analysis around the best paired solution found (see App. G).

We optimize the number of neurons allocated to each type by grid search and their corresponding RFs by gradient descent (Fig. 4, App. G). The four optimal cell type RFs are strikingly similar to those of real ON-OFF midget and parasol cells found in the primate retina (see Fig. 4A-D for representative examples of near-optimal neural network RFs, Fig. 4E-H for macaque data, also see App. I). Both biphasic temporal filters and the characteristic center-surround RF shape are visible. Moreover, consistent with the linear theory, the RF Fourier power spectra of parasol (midget) cells, both in nonlinear simulations and experiments, specialize to cover low (high) spatial and high (low) temporal frequencies. Furthermore, we find that the parasol cells have a higher average firing rate than the midget cells (Fig. 4I), consistent with the greater sensitivity of parasol cells found both in biological data and our linear theory (see Fig. 3C). Also consistent with the linear theory, the neural network optimization loss is reduced for four cell types (two pairs) compared to two (one pair) (Fig. 4J). Moreover, the dependence of performance on cell type ratio mirrors the predictions of our linear theory (compare Fig. 3DE with Fig. 4J and see Appendix H).

**Figure 4:**
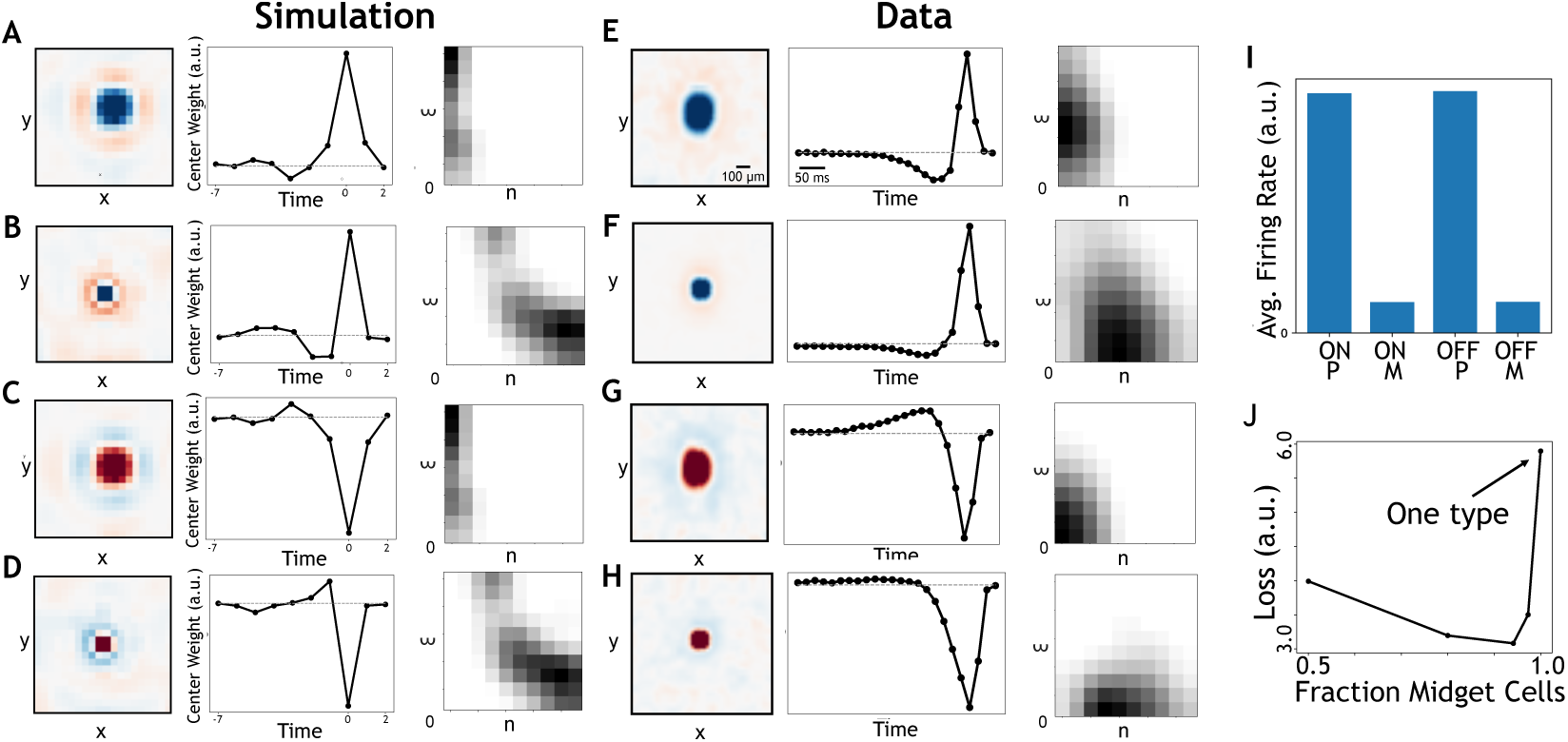
A nonlinear convolutional autoencoder reproduces primate retinal cell types. A, B, C, D) ON parasol-like type, ON midget-like type, OFF parasol-like type, and OFF midget-like type, respectively. Within each panel, far left: spatial receptive field (RF) at peak temporal slice of the spatiotemporal RF. Center: temporal RF, measured as the evolution of the central photoreceptor of the RF across time. Right: Space-time power spectrum of the RF, where dark shades correspond to high power. E, F, G, H) Same quantities measured from macaque retina (see Appendix I). I) Average firing rate per cell of each autoencoder cell type. J) Optimal loss (see Eqn. 3, p=1) as a function of the ratio between cell types densities. Note that optimal reconstruction with two cell type pairs is approximately 2 times better than with one cell type pair (black arrow).

## 6 Discussion

In summary, we first demonstrated mathematically that there is a metabolic advantage to encoding natural movies with more than one convolutional cell type (Fig. 1C). By finding the optimal RFs, strides and cell number ratios for the two populations, we show this advantage is substantial: a 33% reduction in RMS firing rate at a fixed encoding fidelity (reconstruction error: 0.5%). Moreover, the corresponding cell types have similar RFs and densities to midget and parasol cells. We also predict with great accuracy how the ratio of midget to parasol cells varies with the total cell density (Fig. 3F). Finally, by training a nonlinear neural network on the same task of reconstructing natural movies with a limited firing rate budget, we again confirm the advantage of having midget and parasol cells, and we find further differentiation into ON and OFF types.

There are a number of other ganglion cell types found in the primate retina [17] (20 types). Our current model accounts for the four most common cell types (ON and OFF midget and parasol cells), but it could be extended to account for more cell types. The next most common cell type found in the primate retina is the small bistratified type [5], which, unlike midget and parasol cells, pools from blue cones with an opposite polarity to red and green cones. Midget cells are color sensitive [40], a property that we do not account for in our current model, due to our focus on grayscale movies. By taking into account the spatiotemporal statistics of colors in natural movies, one can likely understand the division of labor between midget, parasol and small bistratified cells observed in primates.

Our theory predicts primate cell types well, but interestingly we could not find a good match in other species, such as mouse. The most numerous ganglion cell type in the mouse retina is a selective, non-linear feature detector (W3 cells [41]), thought to serve as an alarm system for overhead predators. Intriguingly, the retina may have evolved to detect behaviorally important predator cues in small animals [7] and efficiently and faithfully encode natural movies in larger animals. A recent study using a deep convolutional model of the visual system suggests that retinal computations either emerge as linear and information preserving encoders, or in the contrary as non-linear feature detectors, depending on the degree of neural resources allocated to downstream visual circuitry [42].

Thus overall our work suggests that the retina has evolved to efficiently encode the translation invariant statistics of natural movies through convolutional operations. Our model strikingly accounts for the 4 dominant cell types comprising 70% of all primate ganglion cells. Furthermore, promising extensions of this work to color statistics could expand the reach of this theory to encompass even greater cell-type diversity.

## Acknowledgements

We thank Alexandra Kling and E.J. Chichilnisky for useful discussions, and for providing us with receptive field visualizations of real midget and parasol cells. We thank Gabriel Mel for a helpful insight about the two-cell proof. We thank the Karel Urbanek Postdoctoral fellowship (S.O) and the NIH Brain Initiative U01-NS094288 (S.D), and the Burroughs-Wellcome, McKnight, James S. McDonnell, and Simons Foundations, and the Office of Naval Research (S.G) for support.

## Appendix

### A Optimizing receptive fields for a dense convolutional array of ganglion cells

In this section, we present the mathematical framework that allows us to find the optimal receptive field for a convolutional array of ganglion cells encoding photoreceptor activations under a constraint on total firing rate [1]. We first solve the problem for one ganglion cell encoding the activation of one photoreceptor in App. A.1. We then use this result to solve for N_C_ ganglion cells of a single type *C* collectively encoding N_p_ photoreceptor activations, where N_*C*_ = N_p_, in App. A.2.

### A.1 Linear reconstruction of a one dimensional signal from a single linear filter

Consider we have a scalar input X drawn from a Gaussian distribution with variance Cxk:

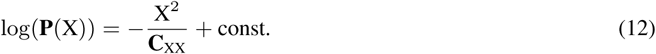

This is measured using a neuron with a scalar filter **F**; this neuron also has intrinsic Gaussian noise *η* with magnitude 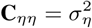, yielding a firing rate of:

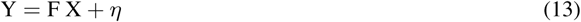

The reconstruction problem is as such: given priors of Eq. 12, which reconstructed signal X_r_ is closest to the actual signal? Because all statistics are Gaussian, this corresponds to finding the most likely X_r_ that produced the observed firing rate, which is given by Bayes rule:

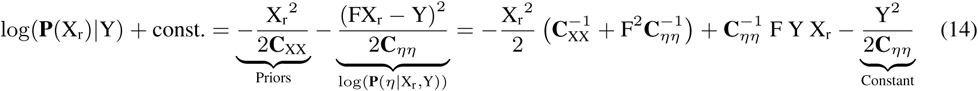

We solve for X_r_ by setting the partial derivative of log probability with respect to X_r_ to be zero:

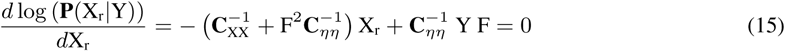

Multiplying each side by C_*ηη*_ C_XX_, we get

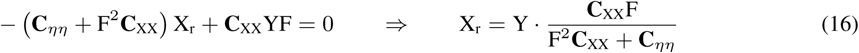

How much variance does the reconstructed signal explain? We note the reconstructed signal is uncorrelated with the reconstruction error, i.e.

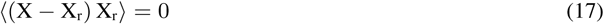

Therefore, the variance explained is simply the variance of the reconstructed signal:

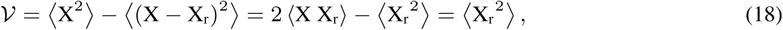

where we have used Eq. 17 in the last step. Using:

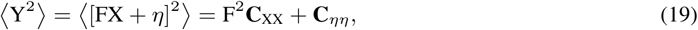

we calculate the variance of the reconstructed signal:

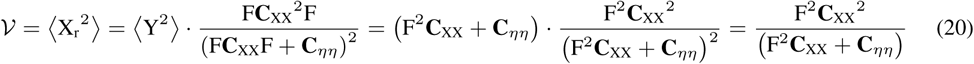

#### Optimal filter magnitude given a penalty on firing rate

Eq. 20 tells us the variance explained as a function of filter strength. Now consider optimizing the filter strength with a penalty term on the variance of firing rate. This corresponds to maximizing the following objective function:

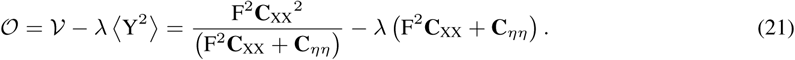

Defining *𝓟* as the power spent encoding the input, *𝓟* = (F^2^C_xx_), this objective can be maximized as:

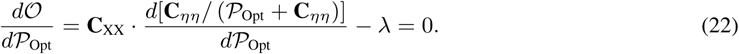

We simplify this as:

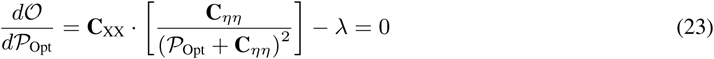

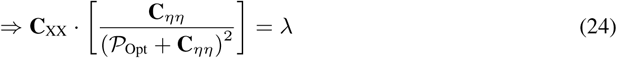

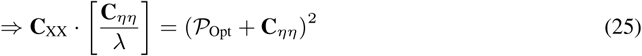

Because 𝓟 ≥ 0, we arrive at our equation for the optimal power spent:

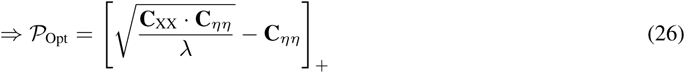

where we use the notation:

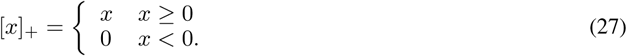

Because C_xx_, C_*ηη*_ are 1 × 1 matrices, we rewrite C_xx_ = S, 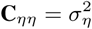. Moreover, because F^2^ = *𝓟*/*S*, we can build similar intuition for the filter magnitude:

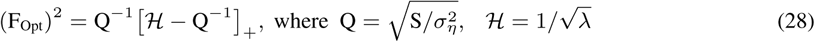

where we call Q the “channel quality” to see that the volume of water (squared filter strength) devoted to encoding corresponds to the volume filled by filling a cylinder of area Q^−1^ and height Q^−1^ to a level of H. The result of Eq. 28 will be used frequently, as the optimization of many convolutional neurons encoding many photoreceptors activations will reduce to it.

### A.2 Convolutional encoding where the number of neurons equals the number of photoreceptors

How do we best encode a photoreceptor image with a convolutional set of linear neurons? We are first going to assume that the number of neurons is equal to the number of photoreceptors, i.e. N_*C*_ = N_p_ = N. In the next section, we then generalize to the case where there can be multiple photoreceptors for each neuron.

Each neuron, indexed by *j*, has a filter F*j* (*i*) *=* F(*i* – *j*) and encodes a photoreceptor image of size N:

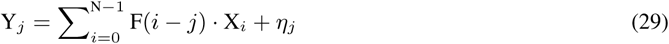

where Y*j* is the neuron activation, X*i* is the activation of photoreceptor *i*, and *η_j_* is the noise at the output of neuron *j*.

It is beneficial to represent the photoreceptor activations, filters, and ganglion cell firing rates in Fourier space. We define the Fourier transform of photoreceptor activations as 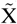, the Fourier transform of ganglion cell activations as 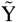, and the scaled Fourier transform of the filter as 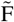:

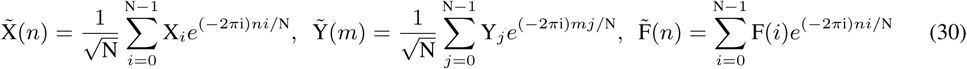

We define 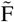 to be the Fourier transform of the filter scaled by 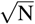 for convenience in future calculations. We can write the signal component of the firing rate of neuron j in terms of the Fourier modes of the photoreceptor activations and filter:

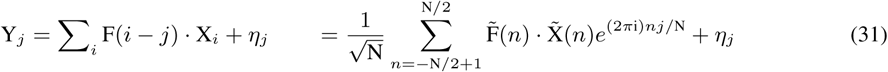

Using this, the Fourier modes of firing rate 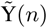 (i.e. the Fourier transform applied on the convolutional array of neurons) can be written as:

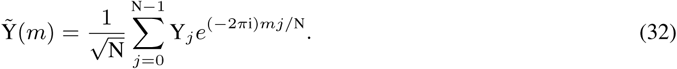

We rewrite the above as:

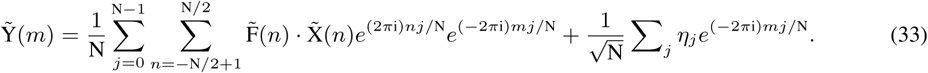

We define 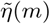 as the Fourier transform of the noise:

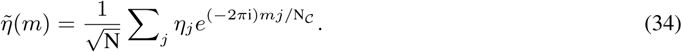

We note that each 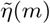 is an independent Gaussian also having variance 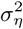 and can rearrange terms:

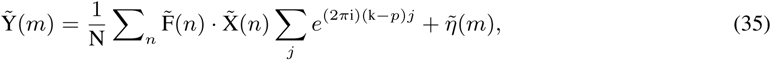

and can use the identity:

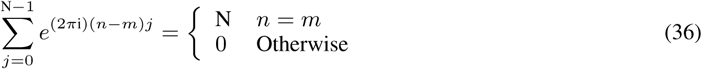

to see that the firing rate at a particular spatial frequency 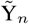 is affected *only* by the photoreceptor image at that same frequency:

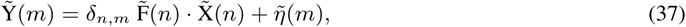

where *δ_n,m_* is the Kronecker delta function. We note that when we add a λ penalty to the total squared firing rate 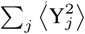, the objective function becomes:

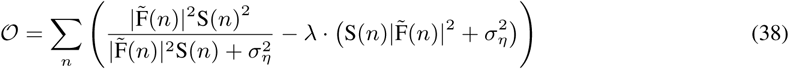

and we see that the objective function Eq. 38 consists of many non-interacting components, each corresponding to a particular spatial frequency *n*, summed together. Moreover, each component corresponding to ganglion cell spatial frequency n is equivalent to the objective maximized in Eq. 21. Therefore, the optimal filters for Eq. 38 are given by Eq. 28:

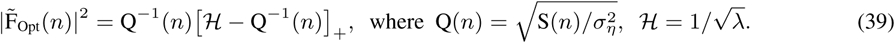

### B Optimal receptive fields for arbitrary ganglion cell density

In App. A, we found the optimal receptive fields for a convolutional array of ganglion cells where the number of ganglion cells was equal to the number of photoreceptors [1]. Now we will solve the more realistic case where the number of ganglion cells is less than the number of photoreceptors. For this, we will first solve the simplified case of one neuron encoding a multidimensional signal (App. B.1), to show that the best single-neuron filter should align with the strongest eigenmode of the covariance. Then, using this result for convolutional encoding with fewer ganglion cells than photoreceptors, we will show that the optimal convolutional filter selects only the strongest N_C_ modes (thus avoiding aliasing [33]), where N_C_ is the number of ganglion cells. The filter strengths are analogous to those solved in App. A.2.

#### B.1 Linear reconstruction of a multidimensional signal from a single linear filter

This section is a generalization of App. A.1 to a single neuron encoding a multidimensional input.

Consider we have a *vector* input 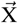 drawn from a Gaussian with covariance C_xx_:

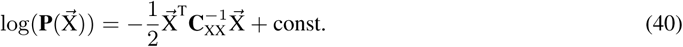

We measure this input using a neuron with receptive field 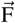 to have a firing rate of:

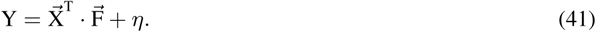

The reconstruction problem is as such: given priors of Eq. 40, solve which reconstructed signal 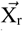 is closest to the actual signal. Because all statistics are Gaussian, this corresponds to finding the most likely 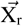 that produced
the observed firing rate:

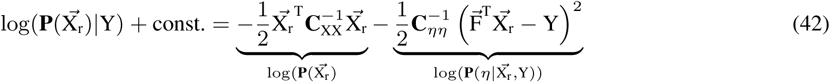

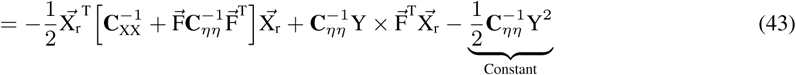

We solve for 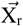 by setting the gradient of log probability with respect to 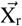 to be zero:

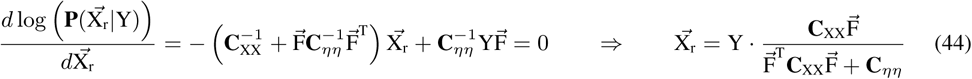

Analogously to App. 18, the magnitude of variance explained is simply the variance of the guess vector 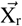. Using:

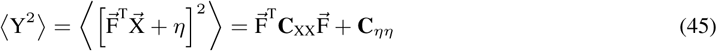

we calculate the variance explained:

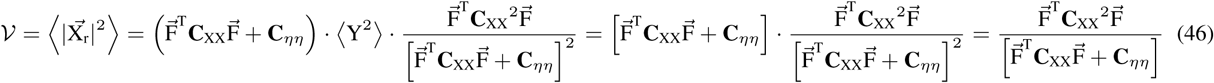

What is the optimal multidimensional filter 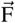 that maximizes variance explained under a penalty on firing rate? Below, we show that this multidimensional problem reduces to a one-dimensional problem by proving that the optimal 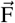 must be a multiple of the largest principal component of C_XX_. Rewriting the 1 × 1 matrix C_*ηη*_ as 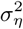, in the eigenspace of Cxk, Eq. 46 can be written as:

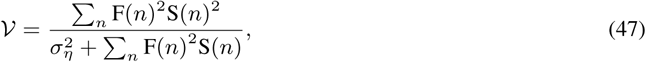

where the eigenvectors of C_xx_ are indexed by *n*, and have eigenvalues S(*n*). Calling the mode index with highest power *n*_o_, we may write the objective as:

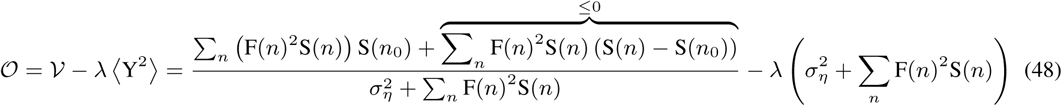

The right term of the numerator can only hurt performance; if there is some other frequency *n* ≠ *n*_0_ with nonzero filters, we note that the transformation redistributing power from mode *n* to *n*_0_:

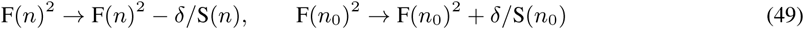

increases the right term of numerator by *δ* · (S(*n*_0_) — S_*n*_) without affecting the total power 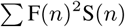, and thus will not affect the rest of the equation. Therefore, the projection should be mapped to only the *largest* mode of the covariance matrix (Fig. 5A), saturating at a performance of:

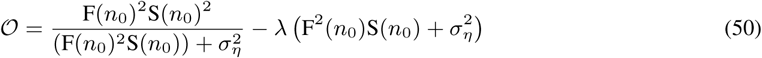

which is optimized in the same manner as the scalar case of App. A.1 to yield

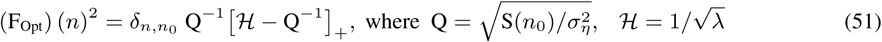

i.e., the 1D optimum after projecting onto the largest eigenmode *n*_0_.

**Figure 5:**
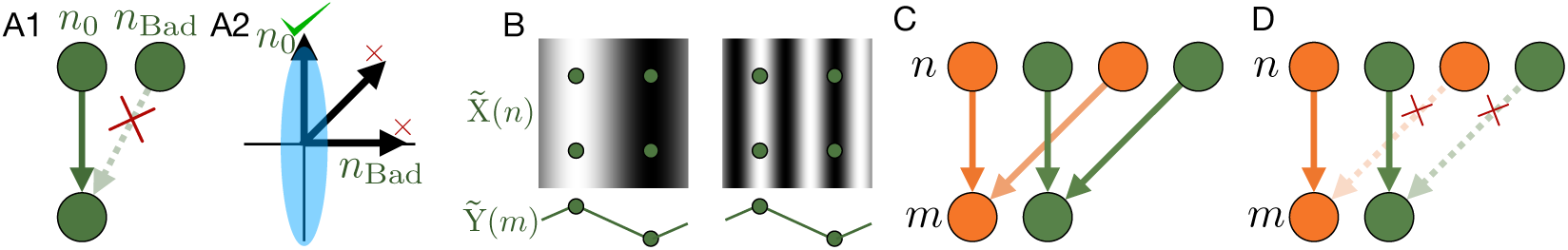
Road map of App. B. A1, A2) When a single neuron can encode multiple inputs (Eq. 41), the optimal filter will be aligned with the largest eigenmode of the input covariance (Eq. 50). B) In our framework, the number of neurons can be smaller than the number of photoreceptors. Therefore, multiple Fourier modes 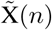 will map onto the same firing modes 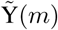. C) Representation of this mapping (Eq. 59). While ganglion cell spatial modes (lower dots) pull from many photoreceptor modes (upper dots), each photoreceptor mode maps to only a *single* ganglion cell spatial mode. D) Therefore, we may reduce this problem to many many non-interacting instances of (A) to show the optimal convolutional filter (Eq. 62) will not be aliased, and each ganglion cell spatial frequency will pull from a single photoreceptor spatial frequency.

#### B.2 Optimal convolutional encoding with fewer ganglion cells than photoreceptors

When there are fewer ganglion cells than photoreceptors, the above framework of App. A.2 must only be slightly modified. Now, instead of one ganglion cell per photoreceptor, we assume a convolutional stride s greater than one, so that each ganglion cell is s apart from its nearest neighbor, yielding a total of N_p_/*s* ganglion cells (we assume N_p_ is a multiple of s). Consider a ganglion cell *j*, centered about *js*, with filter F*j* (*i*) = F(*i* — *sj*). The firing rate of neuron *j* will be:

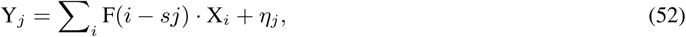

where Y_*j*_ is the neuron activation, X_*i*_ is the activation of photoreceptor *i*, and *η_j_* is the noise at the output of neuron *j*.

It is beneficial to represent the photoreceptor activations, filters, and ganglion cell firing rates in Fourier space. We define the Fourier transform of photoreceptor activations as 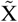, the Fourier transform of ganglion cell activations as 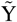, and the scaled Fourier transform of the filter as 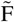:

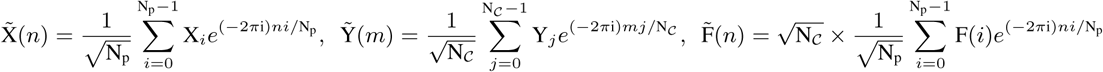

We define 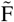 to be the Fourier transform of the filter scaled by 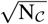 for convenience in future calculations.

We can write the signal component of the firing rate of neuron *j* in terms of the Fourier modes of the photoreceptor activations and filter:

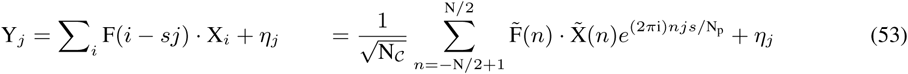

The Fourier modes of firing rate 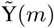 (i.e. the Fourier transform applied on the convolutional array of ganglion cells) can be translated to photoreceptor space:

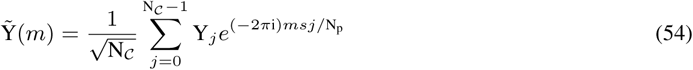

where N_*C*_ = N_p_/s, i.e. the total number of neurons is equal to the number of photoreceptors times the neuron-to-photoreceptor ratio. We rewrite the Fourier transform of firing rates in terms of the Fourier transform of photoreceptor activations and convolutional filter:

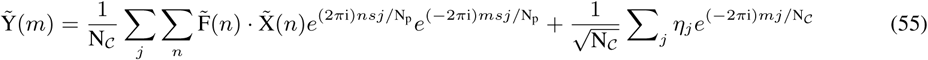

We define 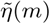 as the Fourier transform of the noise:

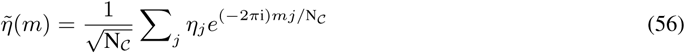

and note that each 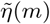 is an independent Gaussian also having variance 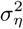. We can rearrange terms:

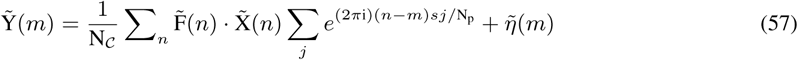

Then, we generalize the identity of Eq. 36 to:

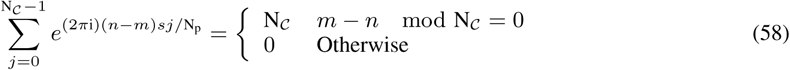

and see that 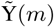 is affected by all 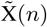 where [*m* – *n*] mod N_*C*_ = 0. This yields the most simplified (Fig. 5C) form of the firing rate in Fourier space:

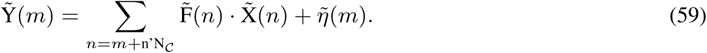

##### Optimal encoding

The objective (Eq. 3) given by this encoding for p = 2 is:

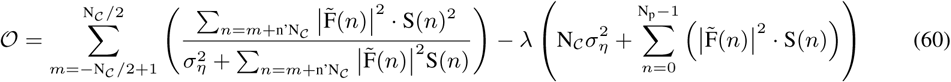

We rewrite Eq. 60 as:

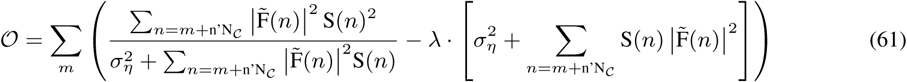

We can verify this step by inspection by noting that each n penalty term is summed over exactly once in both Eq. 60 and Eq. 61.

Examining Eq. 61, we see that the objective function consists of many non-interacting components, each corresponding to a particular ganglion cell spatial frequency *m*, summed together. Moreover, each component corresponding to ganglion cell spatial frequency *m* is equivalent to the objective maximized in Eq. 47. Because, as we saw in App. B.1, it is optimal for a single filter to draw *only* from the largest principal component, the optimal convolutional filter has 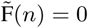 for all *n* ≠ *m* (Fig. 5D) ^3^. Therefore, the optimal filter in frequency space is simply a scaled, truncated version of Eq. 39:

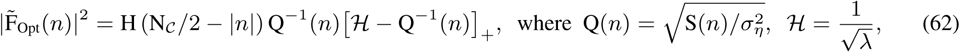

where H is the Heaviside function, which ensures every ganglion cell spatial frequency draws only from a single photoreceptor spatial frequency.

#### C Generalization to encoding of natural movies with spatiotemporal filters

We derived the optimal *spatial* receptive fields for encoding natural *images* in App. B.2. How can we generalize this result to optimal *spatiotemporal* receptive fields for encoding natural *movies?* Now the firing rate of ganglion cell *j* at particular discrete time *t* will be given by:

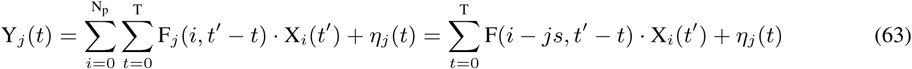

Where T is the length of the movie in frames, and for mathematical convenience we define both the receptive fields and the natural movie to be periodic in space and time.

It is beneficial to represent the photoreceptor activations, filters, and ganglion cell firing rates in Fourier space *and* time. We define the Fourier transform of photoreceptor activations as 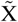, the Fourier transform of ganglion cell activations as 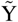, and the scaled Fourier transform of the filter as 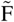:

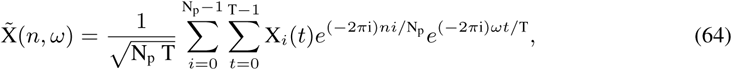

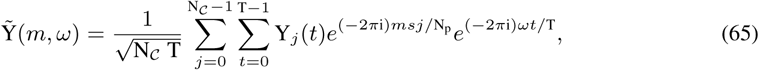

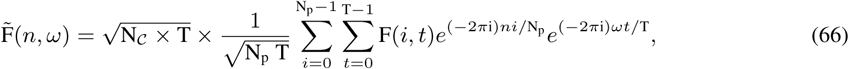

where *t* is an integer representing the frame of the movie, and *ω* is an integer representing the mode number in time. Each *ω* translates into a temporal frequency 2*πω*/T. We define 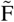 to be the Fourier transform of the filter scaled by 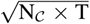 for convenience in future calculations.

This yields the encoding equation in Fourier space:

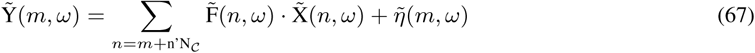

where 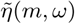 is gaussian with variance 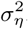. The objective function becomes:

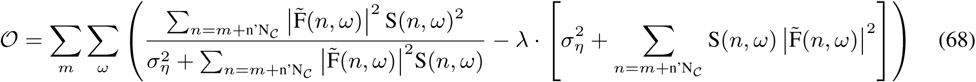

This is nearly identical to the previous case (Eq. 59), except that we have added a time mode. This may be optimized in the same manner^4^ to yield:

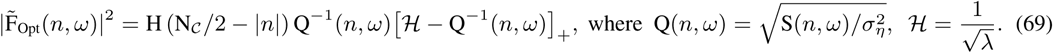

#### D General proof of the benefit of multiple cell types

We will now show that two convolutional types can replicate the encoding fidelity of one convolutional type with less firing rate. For mathematical convenience, we assume an even number of photoreceptors. Given a single convolutional cell type 𝟙 with optimal filters 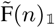, the RMS firing rate (i.e. *p* =1 for Eq. 3) summed over all ganglion cells and all frames in the movie is:

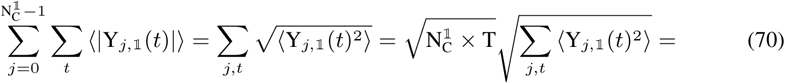

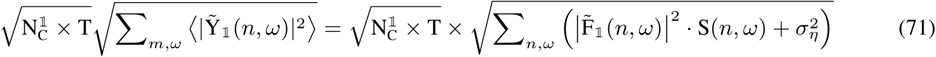

In the main paper and App. E, this constant and the movie length T are factored into the penalty term λ and thus do not appear in the equations.

We can replace one convolutional type with two convolutional types; the first type, 𝔸 covers 𝓝_𝔸_, the first half of all spatial frequencies (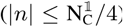), the second type 𝔹 covers 𝓝_𝔹_, the second half of all spatial frequencies (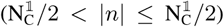). The total number of ganglion cells remains the same, and the stride is doubled: 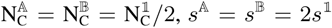. The filters are changed to divide the frequencies between the two cell types.

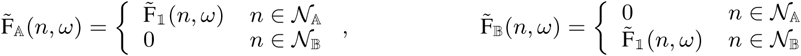

Note that although 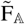, 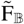 each equal 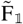, they each have a stronger filter within their respective modes ^5^.

The two cell types presented above have equal variance explained to the single type, as can be seen from the left half of Eq. 68.

When we calculate the total RMS rate of the first cell type:

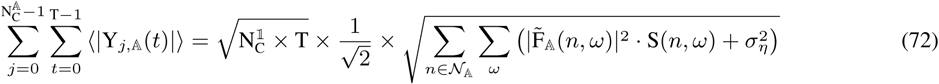

and for the second cell type

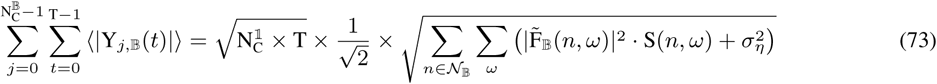

and rewrite the total RMS firing rate for one convolutional cell type as:

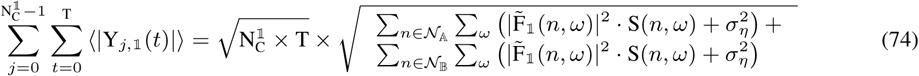

we notice that right square root term for Eq. 74 is simply the sum of the right square root terms for Eq. 72 and Eq. 73. Because the square root of the mean is larger than the mean of the square root, i.e. 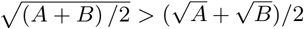 when *A* ≠ *B* (when power monotonically decreases with frequency, A > B), we can show that two convolutional types can achieve *equal* reconstruction error with *less* RMS firing rate. Conversely, they can achieve *better* reconstruction error with the *same* total RMS firing rate.

It is not essential that the mean absolute value of firing rate was penalized. When the penalty on firing rate scales as a power smaller than two, multiple cell types will achieve the same encoding with a smaller penalty, i.e.

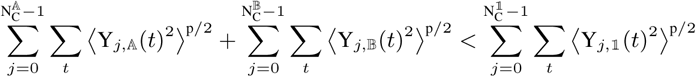

so long as p < 2. We note that this proof construction can be extended to show improvement for arbitrary ratios between the two cell types, as long as (N_*C*_)_A_ + (N_*C*_)_B_ = (N_*C*_)_𝓝_.

##### Many Cell Types

Extending this logic to equally dividing up frequency space among many cell types, the total firing rate becomes:

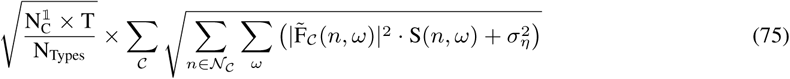

There the total firing of one type likewise can be rewritten as:

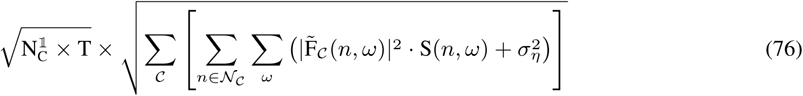

where 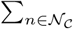 corresponds to the range of spatial frequencies covered by each cell type. We notice that the right square root term in Eq. 76 is simply the sum of the individual right square root terms in Eq. 75. Again, because the square root of the mean is larger than the mean of the square root, many convolutional types achieve the same encoding with a smaller total firing rate.

Because many convolutional types can achieve *equal* reconstruction error with *fewer* total RMS firing rate, they can achieve *better* reconstruction error with the *same* firing rate. In practice, there are diminishing returns with the number of unique cell types.

#### E Objective function and optimization procedure for two cell types

For two cell types, we assume that cells of different types don’t share spatiotemporal modes, yielding an objective function of:

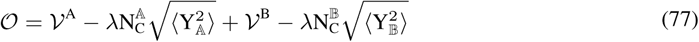

We may replace identically weighted 𝓵_1_ norms on the firing rates of type A and type B cells with differently weighted 𝓵_2_ norms.

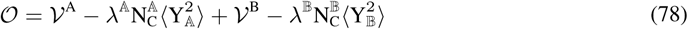

While we know which filters at *fixed strides s*^𝔸^, *s*^𝔹^ and *fixed* 𝗵_2_ *penalties* λ^𝔸^, λ^𝔹^ optimize Eq. 78, we do not know which combination of parameters optimizes these at a *fixed firing budget* (called FR_Max_). To do so, we perform a grid search.

For each combination of *s*^𝔸^, *s*^𝔹^, λ^𝔸^, λ^𝔹^, we optimize the RFs for each type separately. We assign to type A the N_p_/*s*^𝔸^ lower spatial modes and to type B the N_p_/*s*^𝔹^ next modes, as we know that the best encoding scheme will

1. pool only from the modes with highest power and will limit the number of modes selected ^6^ to avoid aliasing,
2. (2) will avoid mixing modes between cell types ^7^, and (3) will favor a solution with a maximal asymmetry between the power of the modes associated to each cell type, so as to maximise the benefit of having two cell types.

We then compute the corresponding reconstruction error and total firing rate. The optimal scheme is the one that minimizes reconstruction error but does not exceed a total firing rate budget ^8^ that we define in advance. We fasten the optimization procedure by working in log space for spatial and temporal frequencies. The code is available online at https://github.com/ganguli-lab/RetinalCellTypes.

Below is a pseudo-code version of the algorithm to maximize variance explained under a fixed RMS firing rate budget of FR_Max_:

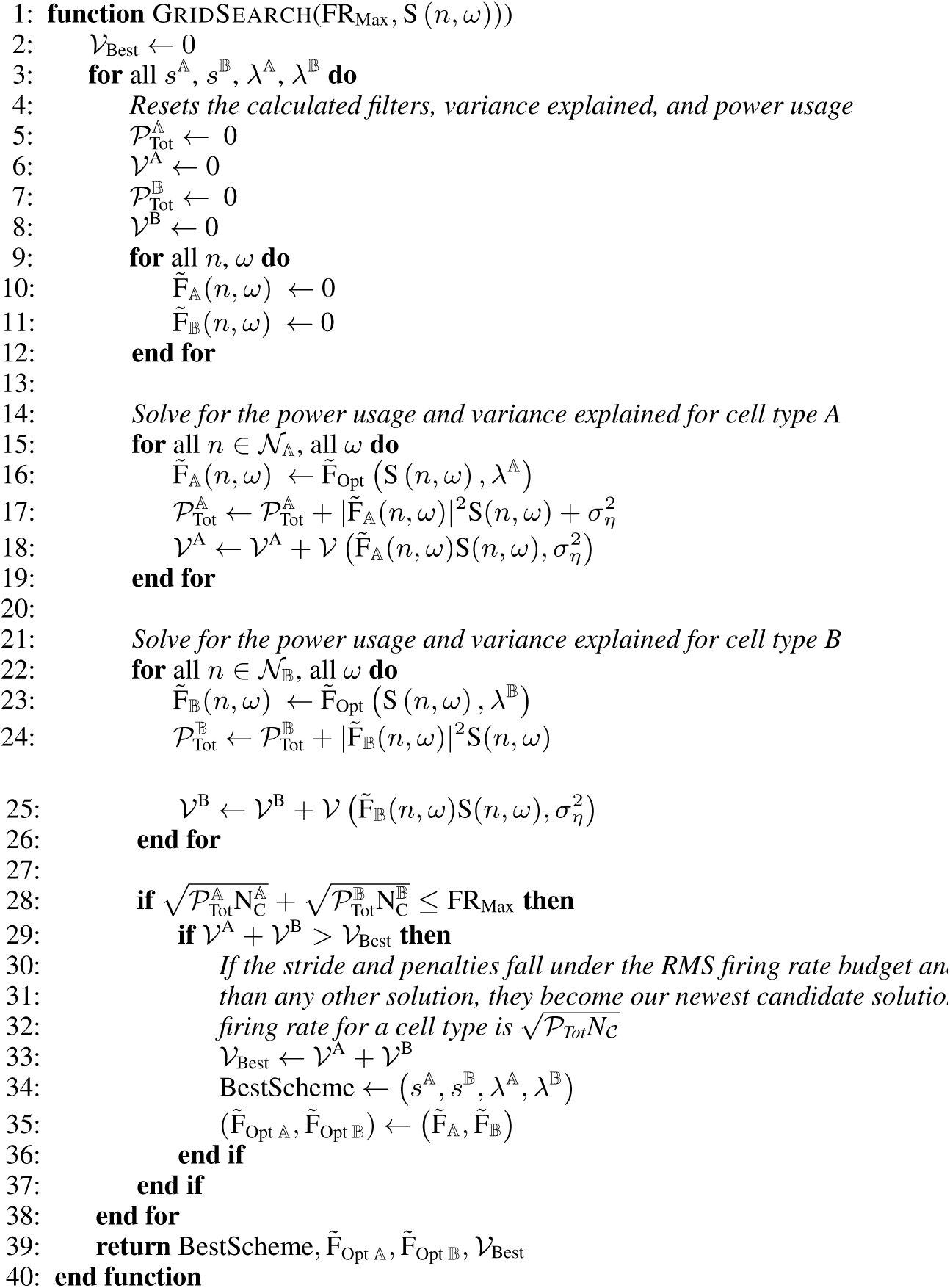
ufig1.

where 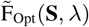 is given by Eq. 39, and where 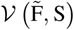 is defined as the total variance explained from that mode in the objective function (Eq. 5):

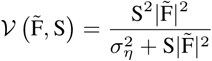

The actual algorithm implemented contains several optimizations for the sake of speeding up the calculation and recycling work, but here we present the “brute force” algorithm for simplicity. We are currently implementing a more sophisticated algorithm that will not necessitate a full grid search (in preparation).

Note that we optimized the two types for the case of one dimension in space and one dimension in time, and we do not consider the contribution of the noise to the mean firing rate to avoid a dependence on temporal frequency cutoffs. We also matched experimental data of Fig. 3F by optimizing cell types with *two* spatial dimensions and time (data not shown, code available online).

#### F Proportion of midget cells as a function of total density in the human retina

To estimate the ratio of midget/parasol cells with respect to total density in the primate retina, we first use the fitted equations of [23] relating the dendritic field size of midget and parasol cells to eccentricty in the human retina:

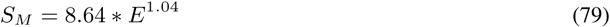

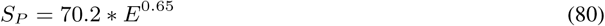

where *S_M_* and *S_P_* are the diameters of the dendritic field of midget and parasol cells respectively and where E is the eccentricity given in mm from the fovea. Then we estimate the corresponding density of midget and parasol cells across retinal eccentricities by using the known overlap between dendritic trees of parasol cells and of midget cells. This overlap has been shown to be constant across the retina, and approximately equal to 1 for midget cells and to 3 for parasol cells [19, 20]. This yields the densities for midget (*D_M_*) and parasol (*D_p_*) cells:

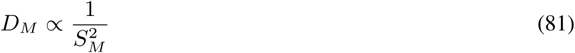

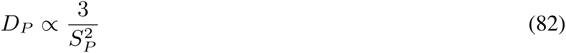

From these equations we can estimate the fraction of midget cells for any total density of cells *D_M_* + *D_p_*. We plot this curve for eccentricities ranging from 0 mm to 20 mm of the fovea.

#### G Neural network simulation

We trained a one-hidden-layer convolutional autoencoder to reconstruct pink noise movies, 10 frames in length with 24×24 photoreceptors, generated according to the spectrum of natural movies. Encoding and decoding layers were implemented as 3D convolutional filters large enough to span the entire movie. A ReLU nonlinearity was applied to the hidden layer and an 𝓵_1_ constraint was imposed on hidden layer activations as in the theory. The generated pink noise dataset contained 50,000 images.

We added a constant noise magnitude of 0.1 to the output of the encoding layer neurons. 𝓵_1_ penalty coefficients of λ = 10^*α*+*β*^ were chosen, with α ∊ {0.5, 0, −0.5} and *β* chosen such that λ (*α* = 0) =4. For a given 𝓵_1_ activation budget, grid search over strides that evenly divide the image width (24 photoreceptors) was used to determine the optimal number of cells (i.e. number of neurons in the hidden layer) for each type (i.e. convolutional channels), and gradient descent was used to optimize receptive fields given these hyperparameters. Each cell type allocation was trained from scratch twice, and plots of performance vs. cell type ratio were obtained by computing the convex lower-bounding hull of all the trials. Cell types were grouped in pairs with an equal number of cells. These were implemented as convolutional channels with strides chosen to yield the desired number of cells. For computational tractability, convolutional filters were given a spatial dilation factor of two. The networks were trained using the ADAM optimizer for two epochs, which was found to yield near-convergence. To bias the network toward respecting causality, convolutional padding was added in asymmetric fashion to the temporal dimension, 70% before the start of the 10-frame movie and 20% after the end, with the total padding sufficient to maintain constant dimensionality in each layer. Thus 80% of encoding and decoding weights connected inputs of earlier time coordinates to outputs of equal or later time coordinates (70% to later coordinates, 20% to equal). We found that learned RFs often contained seemingly extraneous high spatial frequency components; we hypothesize that this phenomenon arises from the fact that high frequency modes have low power and hence, though their presence in the learned filters may not be optimal, they contribute very little to the ¿_1_ firing rate penalty. We show real and Fourier space RFs with Gaussian blur for clarity of visualization.

Note that to make our optimization over four cell types tractable, the grid search over number of neurons per cell type was performed with the restriction that cell types come in pairs with equal numbers of neurons. In fact, primate retinas are thought to contain more OFF ganglion cells than ON cells [23]. This asymmetry has been shown to be an efficient encoding strategy due to the skewness of natural scene statistics, which are not present in our simplified model [13]. Thus, a priori, we had no reason to expect asymmetric ON/OFF population sizes to be beneficial. Nevertheless, we explored the question of whether some asymmetric allocation of cells might be more efficient. We tested the effect of perturbing the cell allocation slightly from its optimal arrangement as determined by the paired-cell-type optimization. Specifically, given the optimal allocation for four cell types of (*N_A_*, *N_A_*,*N_B_*,*N_B_*) cells, we optimized the model for allocations of the form (*N_A_* + *C*, *N_A_* − *C*, *N_B_*,*N_B_*), (*N_A_*, *N_A_*, + *C*,*N_A_* − c), and (*N_A_* + *C*, *N_A_* − *C*, *N_B_* + *C*, *N_B_* − *c*) for small values of *c* = 1,2,3. Thus we perturbed the symmetry of the allocation while preserving the total number of cells. We found that such perturbations reduced the reconstruction performance compared to the optimal model, suggesting that a solution with equal-sized ON and OFF populations of midget and parasol cells is indeed efficient, at least for symmetric Gaussian image statistics.

#### H Effect of firing rate budget on cell type properties

Here we show quantifications of the effect of increasing firing rate budget (or equivalently, decreasing the target reconstruction error). The optimal ratio of midget to parasol cells grows more asymmetric as firing rate penalty decreases in the theory (Fig. 6A) and our neural network simulation (Fig. 6B). Moreover, the benefit of two cell types grows more pronounced as the firing rate budget increases, but is significant across a wide range of parameter settings (Fig. 6C).

**Figure 6:**
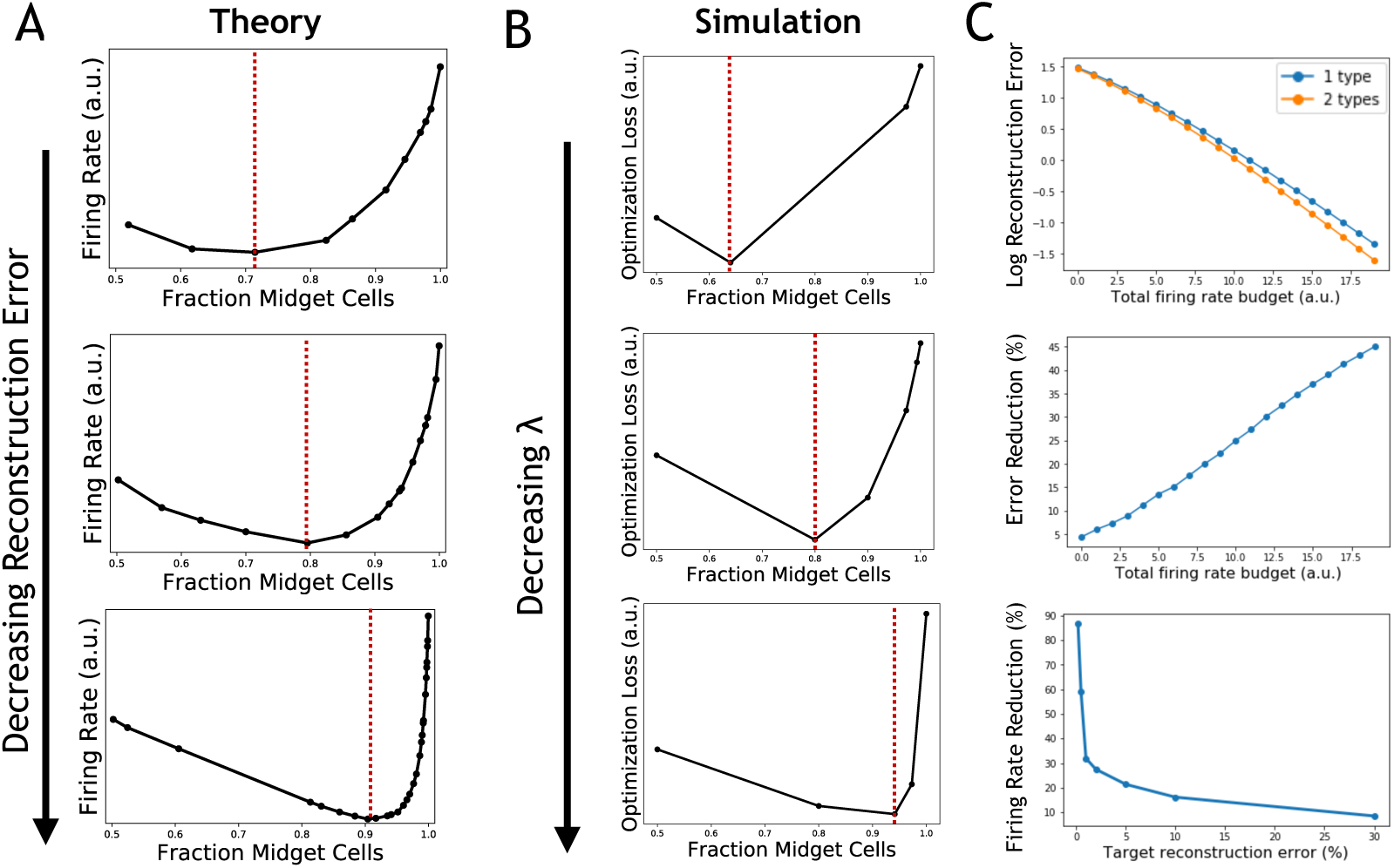
A. Firing rate budget as a function of the fraction of midget cells, for different particular fixed acceptable reconstruction errors (from top to bottom: 25%, 10%, 1%). Note that the optimal cell type ratio shifts to be increasingly asymmetric as the required reconstruction accuracy increases. B. Optimization loss in the neural network simulation as a function of the fraction of midget cells, for different values of λ (from top to bottom, log(λ) = *β* + 1, log(λ) = *β* + 0.5, log(*β*) = *β*, where *β* is a fixed constant). Note that the optimal cell type ratio shifts to be increasingly asymmetric as the required reconstruction accuracy increases. C. Top: Reconstruction error as a function of firing rate budget, for optimal one cell type and two cell-type solutions. Error decreases as budget increases, two cell types outperform one, and the disparity grows with the firing rate budget. Middle: The relative reduction in error at varying firing rate budgets for the optimal two cell type solution as compared to the optimal one cell type solution. Bottom: The relative reduction in firing rate required to achieve varying target reconstruction errors for the optimal two cell type solution as compared to the optimal one cell type solution. Note that the benefit is significant across a wide range of target reconstruction errors, and it is especially high when high accuracy is required.

#### I RFs of real midget and parasol cells

Receptive fields (RFs) of real midget and parasol cells were obtained by measuring ganglion cell responses of an explanted macaque retina with a multi-electrode array [43], in response to a randomly flickering black&white checkerboard stimulus. The checker size was 22.5 micron on the retina; the frame rate was 60 Hz. After isolating cells with a spike sorting procedure [44] and identifying ON and OFF midget and parasol cells [43], the spike-triggered-average (STA) of each cell was computed, and an average RF was obtained for each cell type by aligning each cell RF of a given type with respect to their center (defined as the max intensity pixel of the RF of each cell), and then applying a smoothing function on the average RF (default bicubic function of MATLAB - weighted average of pixels in the nearest 4-by-4 neighborhood).

We then performed the same analyses on these RFs that we did for our simulated RFs in Fig. 4.

Full spatio-temporal RFs for simulated and real ganglion cells are shown in Fig. 7.

**Figure 7:**
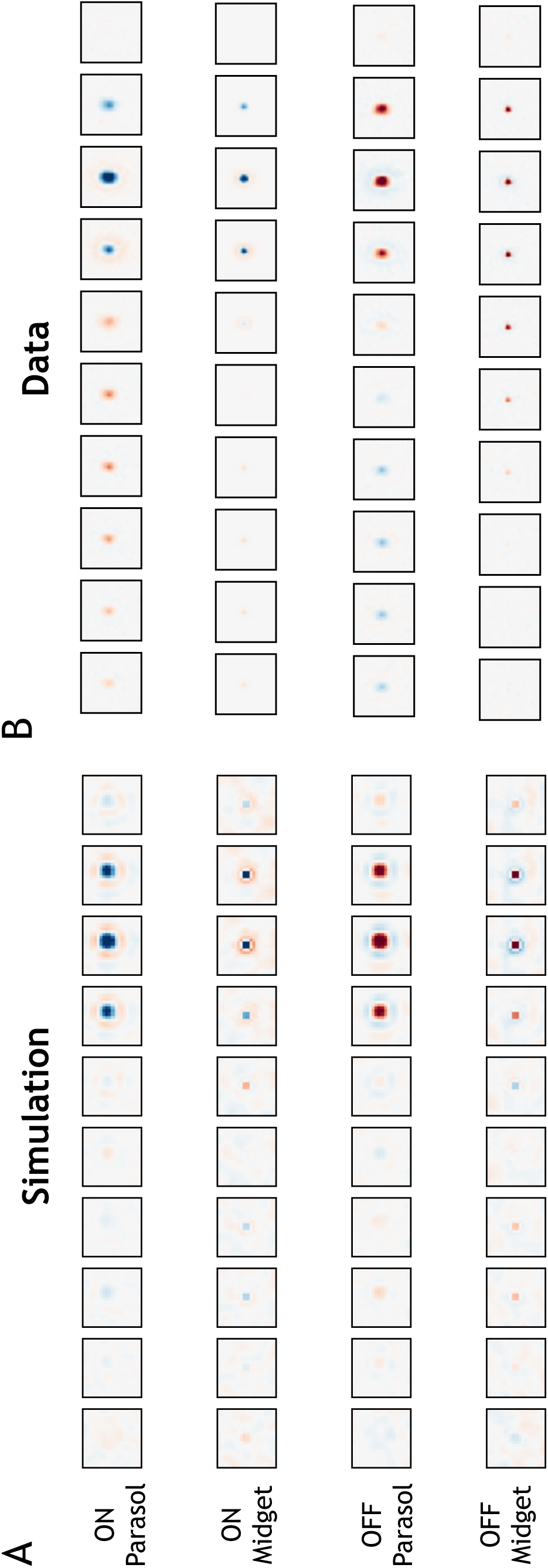
A. Receptive fields for four cell types across time, as computed by the neural network simulation (same settings as in Figure 4). B: Spike-triggered averages for real macaque cells. One frame corresponds to a temporal bin of 16.7ms.

## Footnotes

3 3Because the power decreases monotonically with frequency, the highest power image frequency corresponding to *m* is simply given by *n*M(*m*) = *m*.

4 The optimum could have each ganglion cell spatial frequency pull from multiple photoreceptor spatial frequencies, where the spatial frequency pulled from was a function of the temporal frequency. This will not happen for natural movie statistics, since the power distribution monotonically decreases with both spatial and temporal frequency

5 This is because 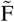 is the Fourier transform of the filter strength *scaled by the square root of the number of neurons*. The magnitude of the Fourier transform of F_𝔸_, F_𝔹_ have each increased by 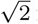 for their respective modes. The intuition behind this is that when encoding a mode at fixed SNR with a variable number of neurons,the *total* power devoted to encoding that mode must remain constant. Therefore, when there are fewer neurons, each neuron must dedicate more power, and thus a stronger filter, to that mode

6 6The number of modes selected will be equal to the number of cells of that type (see App. B.2).

7 7This grid search assumes that in the optimum, there is no mode *n* for which both 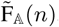,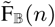 are nonzero, i.e. no mode is shared between the two cell types. We find this to be empirically true from optimizing the linear problem numerically with gradient descent (code available online).

8 We also use a variant which minimizes total firing rate without going below a minimal coding fidelity

